# Treg fitness as a biomarker for disease activity in Juvenile Idiopathic Arthritis

**DOI:** 10.1101/2024.04.24.590917

**Authors:** Meryl H. Attrill, Diana Shinko, Telma Martins Viveiros, Martina Milighetti, Nina M. de Gruijter, Bethany Jebson, Melissa Kartawinata, Elizabeth C. Rosser, Lucy R. Wedderburn, CHARMS study, JIAP study, Anne M. Pesenacker

## Abstract

Juvenile Idiopathic Arthritis (JIA) is an autoimmune condition characterised by persistent flares of joint inflammation. However, no reliable biomarker exists to predict the erratic disease course. Normally, regulatory T cells (Tregs) maintain immune tolerance, with altered Tregs associated with autoimmunity. Treg signatures have shown promise in monitoring other autoimmune conditions, therefore a Treg gene and/or protein signature could offer novel biomarker potential for predicting disease activity in JIA.

Machine learning on our nanoString Treg gene signature on peripheral blood (PB) Tregs generated a model to distinguish active JIA (active joint count, AJC≥1) Tregs from healthy controls (HC, AUC=0.9875). Biomarker scores from this model successfully differentiated inactive (AJC=0) from active JIA PB Tregs. Moreover, scores correlated with clinical activity scores (cJADAS), and discriminated subclinical disease (AJC=0, cJADAS≥0.5) from remission (AUC=0.8980, Sens=0.8571, Spec= 0.8571).

To investigate altered Treg fitness in JIA by protein expression, we utilised spectral flow cytometry and unbiased analysis. Three Treg clusters were increased in active JIA PB, including CD226_high_CD25_low_ effector-like Tregs and CD39-TNFR2-Helios_high_, while a 4-1BB_low_TIGIT_low_ID2_intermediate_ Treg cluster predominated in inactive JIA PB (AJC=0). The ratio of these Treg clusters correlated to cJADAS, and higher ratios could predict inactive individuals that flared by 6-month follow-up.

Thus, we demonstrate altered Treg signatures and subsets as an important factor, and useful biomarker, for disease progression versus remission in JIA, revealing genes and proteins important in Treg fitness. Ultimately, PB Treg fitness measures could serve as routine biomarkers to guide disease and treatment management to sustain remission in JIA.

## Introduction

Juvenile Idiopathic Arthritis (JIA) is the most common rheumatic autoimmune condition with childhood onset. Persistent, uncontrolled inflammatory flares of joints lead to pain, reduced quality of life, and disability(1). Targeted biologics, such as TNF-α blockade, and more general immunosuppression by corticosteroids and methotrexate have improved disease management, yet 30-50% of patients do not achieve adequate responses(1, 2). Furthermore, these therapeutics can often cause severe side effects(3) and with no clear guidelines on when to withdraw safely, patients may remain on medication for longer than necessary due to a high risk of flare(4). Treatment decisions are often based on clinician’s experience and preference, with limited objective quantifiable markers to measure disease activity(1, 4). Predicting flares, monitoring response to treatment, and identifying true remission from subclinical disease therefore remain key challenges in JIA to achieve sustained remission(1, 5).

Clinically applicable biomarkers are valuable tools in governing treatment decisions and measuring clinical outcomes. Routinely measured biomarkers in JIA include autoantibody/rheumatoid factor (RF), determining subtypes of polyarticular JIA, and anti-nuclear antibody (ANA) which associates with increased risk of developing uveitis(5). However, these markers are utilised for classifying disease subcategory and risks of co-morbidities, rather than monitoring disease progression and predicting flares. Cellular ratios, gene expression profiling, and proteome analysis of synovial fluid mononuclear cells (SFMCs) have been suggested to identify JIA individuals likely to progress into a more severe category of disease activity to extended oligoarticular JIA(6, 7). However, SFMCs are only present and accessible during an active inflammatory flare of the joint, and therefore are not a viable source to measure subclinical disease. In blood, an RNA signature of 99 genes was able to segregate JIA patients which achieved remission on methotrexate from non-responders, using blood samples prior to treatment commencing(8). However, large genetic signatures and complex techniques are not suitable for a standardised, clinically-applicable biomarker(5). Increased serum concentrations of pro-inflammatory calcium-binding S100 proteins, such as S100A8/A9 and S100A12, have predicted response to methotrexate and anti-TNF therapy(9–11) and can assist with assessing the risk of flare and guide timing of MTX withdrawal in clinical remission(12, 13). Although now adopted in some clinical centres as subclinical inflammatory markers to assist therapeutic decisions in JIA, these S100 proteins have shown more efficacy in monitoring systemic disease(14), can lack biomarker specificity by misclassifying a large percentage of patients(10), and are not yet standardised. Therefore, no reliable, validated biomarker in JIA currently exists to predict the erratic disease course, determine which individuals are at risk of imminent flares, or when to taper medication off safely once sustained remission is achieved.

Regulatory T cells (Tregs) are key in the control of immune homeostasis and usually prevent inappropriate inflammation through maintaining tolerance. When Tregs fail to control effector cells, autoimmunity, such as JIA, can arise. Tregs have therefore been investigated for their therapeutic potential in several other autoimmune conditions(15). Treg dysfunction is also a hallmark of JIA(1, 16, 17), yet immune regulation has yet to be targeted or monitored for therapeutic benefit in JIA. CD4+Foxp3+ Treg changes at the site of inflammation in JIA have been well described, with enrichment in synovial fluid (SF) and large heterogeneity(18–20), suggesting that the presence of specific Treg subsets, with unique co-receptor combination, likely have different functions. Interestingly, SF Tregs retain hypomethylation of the TSDR (Treg specific demethylated region), showing commitment to the Treg fate(21) and increased numbers may associate with less extensive disease(22). However, several genetic risk alleles in JIA are in loci associated with Treg function(16), with synovial Tregs suggested to exhibit a pro-inflammatory effector phenotype, loss of IL-2 sensitivity and questioned *in vitro* suppressive capacity(17, 18, 21, 23–27). Investigations into altered Treg phenotype in the blood between inactive and active JIA are more limited in non-systemic JIA.

In other conditions, Treg abundance and Treg markers have been investigated as predictive biomarkers for disease manifestations, therapeutic response, and prognosis, in cancers(28), cardiovascular disease(29, 30) and autoimmunity(17, 31, 32). Although large gene signatures are impractical for standard clinical use, nanoString technology requires comparatively few cells from a sample (∼5000 cells), with no RNA purification, no amplification bias and high replicability(17, 33, 34). A nanoString signature as a biomarker would therefore be feasible for clinical use, having already been approved clinical use for cancer prognosis(35).

We have previously developed a nanoString RNA Treg signature, incorporating genes which reflect Treg function, genes consistently expressed by healthy Tregs, with so far undefined functions, and effector-linked genes, thus discriminating Tregs from conventional T cells (Tconv) regardless of activation state(31). On purified Tregs, the Treg signature assesses intrinsic Treg changes(31, 32), thus infers functionality of Tregs. Indeed, this signature sensitively and specifically identified children and adults with new onset type 1 diabetes (T1D) from healthy controls, as well as predicting responders to biologic treatment in T1D patients and disease trajectory(31, 32). With genetic similarities to T1D, including key genes involved in Treg functions(36, 37), here we further investigate our Treg signature in oligoarticular and RF-polyarticular JIA, incorporating 11 additional genes of pathways previously linked to JIA and/or autoimmunity (48 gene Treg signature Plus, supplementary Table S1), and utilise high dimensional spectral flow cytometry data to asses Treg fitness profile at protein level.

We explore the biomarker potential of Treg fitness signatures in differentiating inactive from active JIA blood, identifying subclinical disease, and predicting disease progression. By monitoring changes in Tregs through mRNA and protein as reflective measure for *in vivo* Treg fitness in the periphery in JIA, it may be possible to identify disruption to the immunoregulatory balance prior to cellular infiltration and inflammation of the joint. Therefore, therapeutic preventatives can be introduced earlier to sustain remission, or medication withdrawn safely when remission is indicated.

## Results

Here we focus on the most common subtypes of JIA: oligoarticular (persistent and extended) and RF negative polyarticular JIA, as they are largely similar in immune profile(38), and utilise active joints as a main classifying criterion of disease activity. To assess biomarker potential of Treg fitness signatures in JIA, mononuclear cells (MCs) and Tregs (purity sorted CD4+CD25highCD127low) were isolated from peripheral blood (PBMC n=59, Treg n=54 post data QC) and unmatched synovial fluid (SFMC n=28, SF Treg n=26 post data QC) of active joints from individuals with JIA. PBMCs and Tregs from adult healthy controls (aHC, PBMC n=20, Treg n=24 post data QC) and paediatric healthy controls (pHC, PBMC n=5, Treg n=3 post data QC) were additionally included as separate groups.

### 48 gene Treg signature can discriminate PB Tregs from unfractionated cells and SF Tregs

Multivariate principal component analysis (PCA) was performed on normalised, log_2_-transformed Treg signature Plus (supplementary Table S1) nCounter data from Tregs and unfractionated cells of all groups (Figure 1). The first two principal components (PC) represented 62.4% of the total variance of the eight sample groups. As expected, PC1, explaining 47.6% of the variance, categorically separated Tregs from unfractionated cells from the same sample group (Figure 1), demonstrating the Treg-specific combination of genes in the signature. Tregs and MCs from SF additionally clustered discretely from those from PB, represented by PC2, explaining 14.8% of the variance (Figure 1). Whilst PBMCs from aHC, pHC, and individuals with JIA had overlapping clusters in PC2, PB Tregs showed greater divergence between these groups (Figure 1). Interestingly, JIA PB Tregs, taken together from clinically active and inactive JIA patients, defined by active joint count (AJC≥1 and AJC=0 respectively at time of sample), spanned the largest range in PC2 of all groups (Figure 1). PB Tregs could therefore be effectively discriminated from unfractionated cells and Tregs from the inflamed joint using our 48 Treg signature Plus gene set. This further emphasises possible altered functional properties of synovial Tregs at the site of inflammation in JIA (18–20), and large variability in PC2 of JIA PB Tregs suggests possible heterogeneity in the blood.

**Figure 1.**
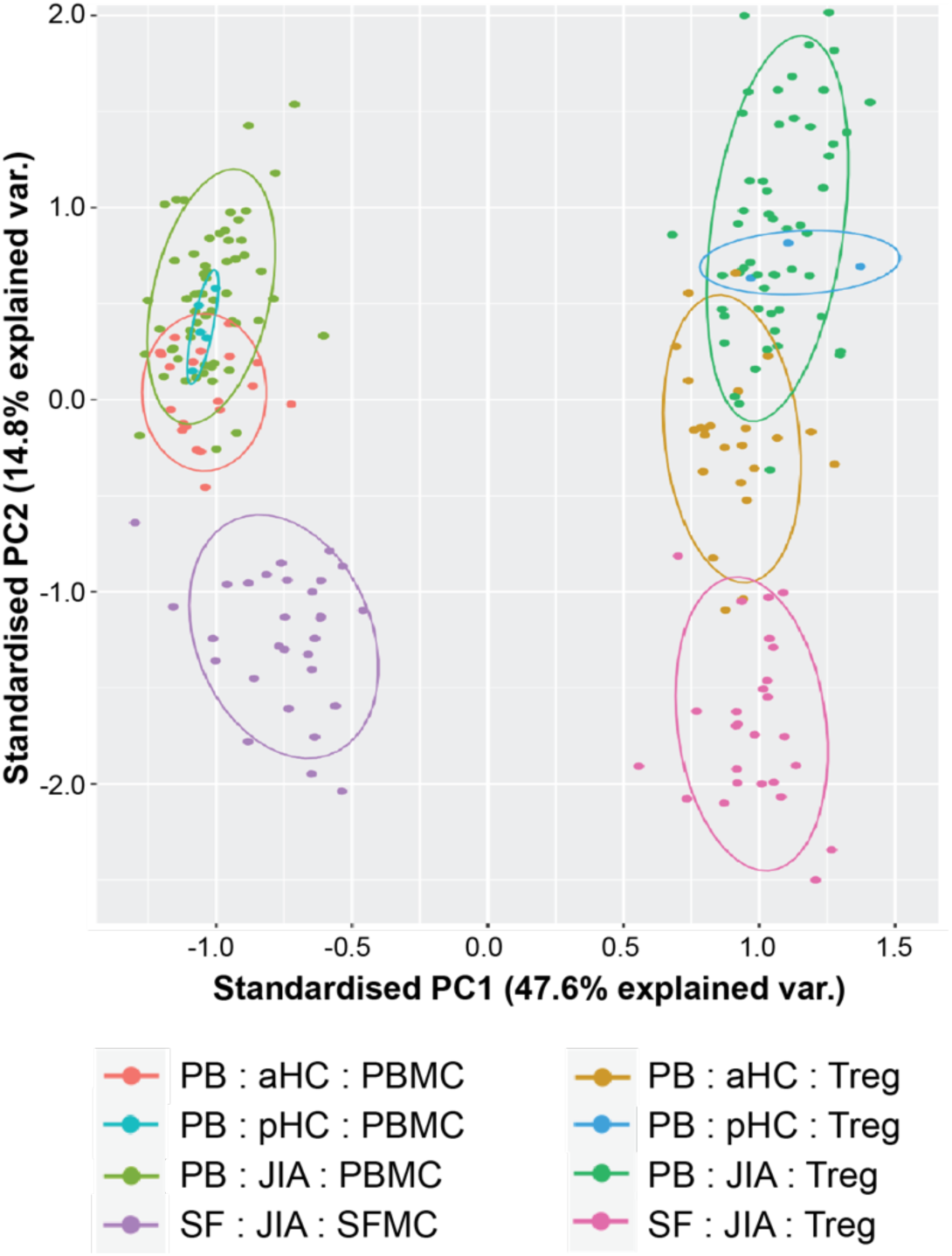
Principal component analysis (PCA) of Tregs and unfractionated cells in HC and JIA using 48 gene Treg signature. PCA performed on normalised log2-transformed nanoString data (mRNA count) for 48 Treg signature Plus genes. First two principal components, explaining 62% of total variance displayed with corresponding percentage. Circles represent normalised data points with ellipse at 0.8 confidence interval. Tregs (sorted CD4+CD25highCD127low) and unfractionated cells were paired where available. aHC= adult healthy control; pHC= paediatric healthy control; JIA= Juvenile Idiopathic Arthritis; PB= peripheral blood; SF= synovial fluid; MC= mononuclear cell.

To further investigate the variance found in PC2, and whether changes in Treg fitness at the site of active inflammation in JIA are reflected by altered Treg signature in PB, we utilised machine learning to develop models to predict active disease from sorted Treg gene signatures. Genes which had no detected counts for more than 30% of all samples in all groups were removed from further analysis and samples were split into equally sized train and test sets. As pHC PB samples were limited, aHC PB Tregs were used in training sets for healthy controls, with pHC PB Tregs ran in separate test sets for age-matched validation.

### Synovial Tregs have an altered RNA profile which is not reflected in the periphery of active disease

Firstly, an elastic net regression generated a model from 42 normalised, scaled mRNA counts from the Treg signature (Figure 2) producing SF Treg signature scores which perfectly distinguished aHC PB Tregs from JIA SF Tregs (AUC= 1.000, Sens= 1.000, Spec= 1.00, p<0.0001, Figure 2B), aligning with previous findings of functionally distinct Tregs in the inflamed environment(18–20). Genes with the greatest coefficient weighting in the model (Supplementary Table S2) included inflammatory cytokine receptor IL1R1, genes important for translational processes (EIF3S6, involved in translation initiation; HNRPA1 in pre-mRNA processing and transport, and RPL23A, a ribosomal protein), and key Treg-associated genes CTLA4 (CD152), FOXP3, TIGIT, and TNFRSF1B (TNFR2/CD120b) (Figure 2A). This suggests a translational reprogramming of synovial Tregs in the inflamed environment and altered expression of key markers involved in Treg fitness.

**Figure 2.**
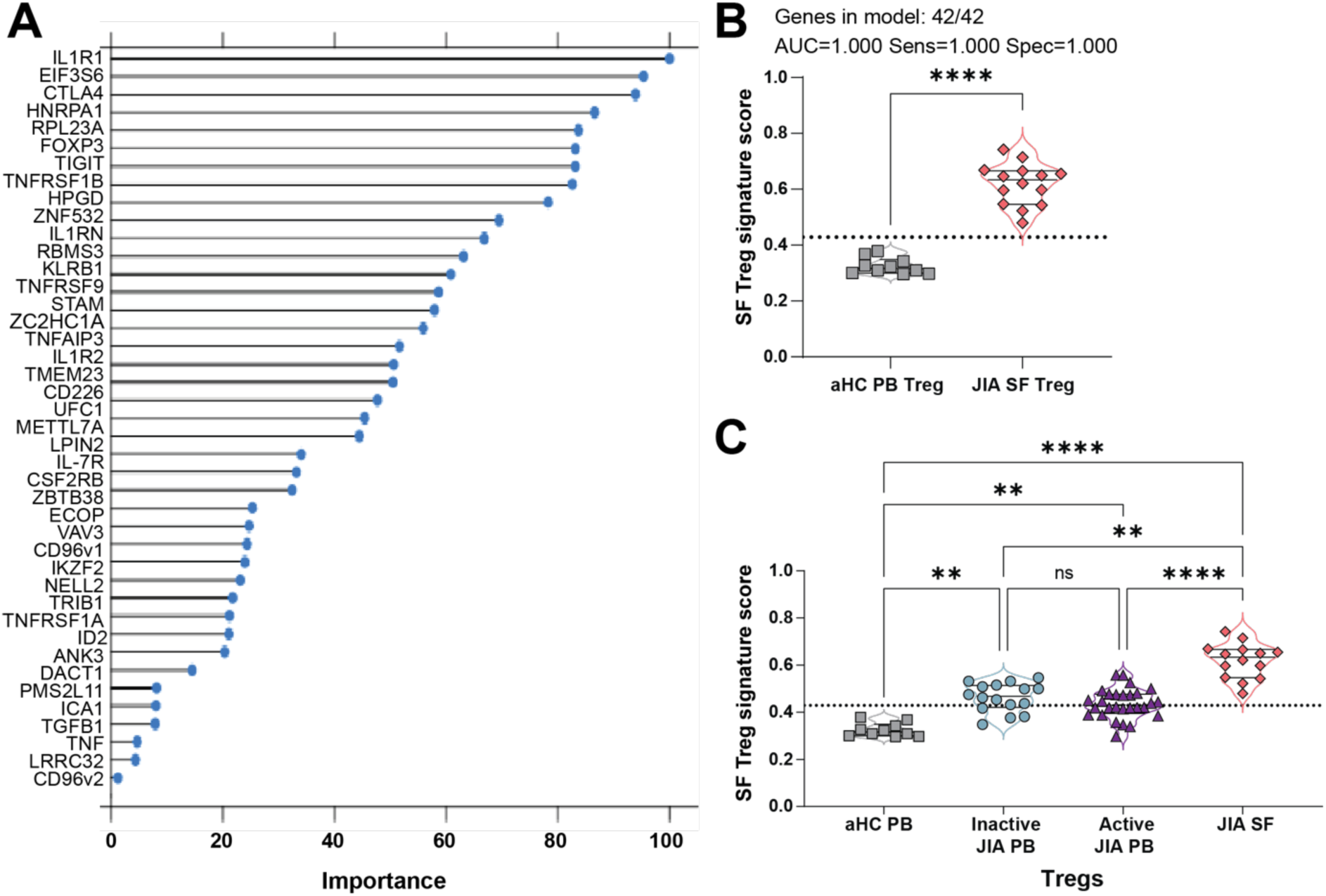
SF Treg signature scores can distinguish blood from synovial Tregs but does not relate to disease activity in peripheral blood of JIA. Treg signature Plus mRNA counts by nanoString were normalised, log2-transformed, and split into 50/50 train and test sets before scaling. Utilising our biomarker discovery pipeline, optimum parameters were chosen by elastic net regression on 42 Treg signature genes with leave one out cross validation (LOOCV) for best accuracy in classifying Tregs (sorted CD4+CD25highCD127low) from adult healthy control PB (aHC, signature score assigned towards 0) and JIA SF (signature score assigned towards 1). **A)** Importance of the 42 genes in model selection, ranked by frequency of each gene used to differentiate groups in each iteration (importance as %). **B)** SF Treg signature score of test set, with 0 being most like aHC PB and 1 being most like JIA SF from the model derived through biomarker discovery pipeline. AUC, sensitivity, and specificity shown at cut off 0.4290 **C)** Inactive JIA (AJC=0) PB Tregs and active (AJC≥1) JIA PB Tregs ran as additional test dataset on this model, with test SF and aHC SF Treg signature scores displayed. For B-C, Mean ±SEM displayed, with individual points; cut offs for Sens/Spec calculation displayed by dotted line; Significance determined by Mann-Whitney test (B), and one-way ANOVA with Kruskal-Wallis multiple comparisons post-hoc test (C) with **p<0.01, ****p<0.0001, ns= not significant. PB= peripheral blood; SF= synovial fluid; JIA= Juvenile Idiopathic Arthritis; AJC= active joint count; AUC= area under curve; Sens= sensitivity; Spec= specificity.

To determine whether this specific SF Treg signature could be reflected in the periphery of active JIA, we ran the same model on a test set of JIA PB Tregs from individuals with one or more actively inflamed joints (AJC≥1, n=28) at the time of sample, and from clinically inactive (no active joints, AJC=0, n=16). Generated SF Treg signature scores therefore reflected the probability of a sample being classified as most similar to SF Tregs (SF Treg signature score=1) or PB Tregs from aHC (Treg SF signature score=0). In accordance with the PCA analysis (Figure 1), both inactive and active JIA PB SF Treg signature scores were significantly different from JIA SF Tregs (p=0.0077 and p<0.0001 respectively) and aHC Tregs (p=0.0016 and p=0.0096 respectively) (Figure 2C). However, no difference was found in SF Treg signature scores between inactive and active JIA PB Tregs (p>0.9999, Figure 2C).

We therefore conclude that SF-derived Treg signatures are not viable biomarkers in distinguishing active from inactive JIA in blood due to significant adaptions SF Tregs undergo within the inflamed joint.

### Blood Treg-derived signature scores can distinguish active from inactive disease in JIA and identify subclinical disease activity

Next, we focused on blood-derived models to differentiate disease activity in JIA. As the number of inactive JIA PB samples acquired were too limited to allow for train and test set splits, we trained a model to distinguish aHC Tregs from active (AJC≥1) JIA PB Tregs (Figure 3). Optimal elastic net regularisation parameters were chosen by leave one out cross-validation, leading to a final model incorporating 23 out of 37 genes (supplementary Table S2) which successfully differentiated the test set aHC PB Tregs from JIA PB Tregs (AUC=0.9875) with perfect specificity (1.000) and high sensitivity (0.9375), at a Treg signature score cut off of 0.5 (Figure 3A,B). The top three genes of most importance in biomarker discovery pipeline, ZNF532, TNFRSF9 (CD137/4-1BB), and IL7R (CD127), are likely to be age-related, as demonstrated in the mean expression between groups (Figure 3D, supplementary Figure S1). Nonetheless, Tregs from paediatric HC (pHC) PB input as a separate test set were significantly different in signature scores to active JIA PB (p=0.0144, AUC=0.9375, sens=0.9375, spec=0.6667 at cut off 0.5, Figure 3C), suggesting that although age may be a factor, most genes governing this model are disease-dependent. Key genes in the model which distinguished Tregs of individuals with active JIA from healthy controls included: ZC2HC1A (C8ORF70), CSF2RB, UFC1, CD96v2, TNFRSF1B (CD120b/TNFR2), STAM, HNRPA1, TRIB1, CTLA4 (CD152), HPGD, and TNFAIP3 (A20) (Figure 3A,D). While most genes on their own did not show a statistically significant difference in expression between active and inactive PB Tregs, CTLA4 mRNA was slightly, but significantly, increased in Tregs from active JIA PB (mean±SEM log_2_-transformed mRNA count inactive 8.354±0.10 vs active 8.587±0.07, p=0.0432, Figure 3E), potentially indicating a heightened activation state(39). Interestingly, the co-receptor isoform CD96 variant 2 (CD96v2) mRNA was significantly decreased in PB Tregs from patients with active joint inflammation (inactive 4.243±0.32 vs active 3.278±0.30, p=0.0040, Figure 3E). CD96 competes with TIGIT and CD226 for ligand CD155, with CD96v2 showing higher binding capacity compared to variant 1 isoform(40). However, the role of CD96 in Treg fitness remains unclear.

**Figure 3.**
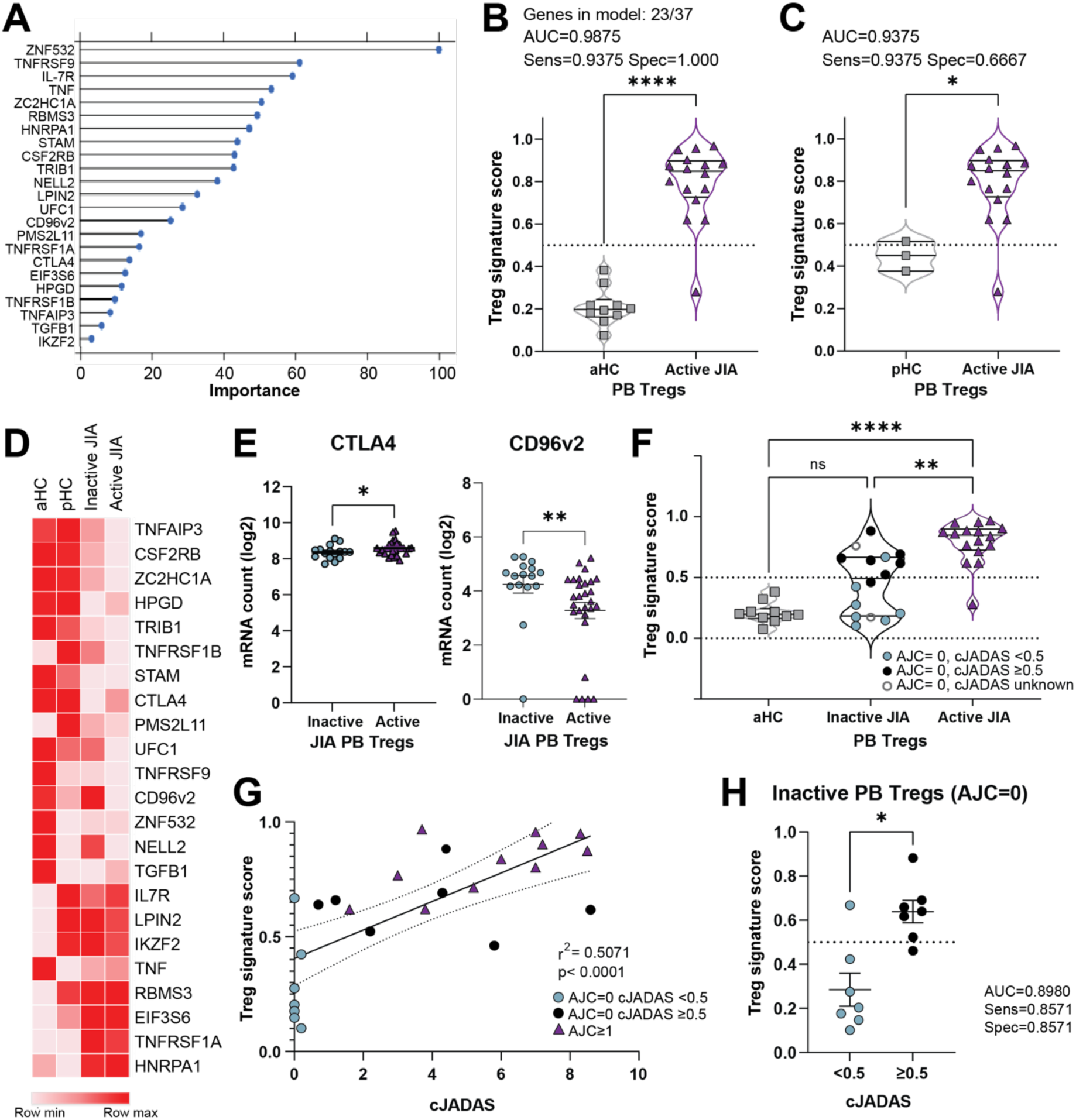
PB-derived Treg signature successfully differentiates active from inactive JIA, correlates to disease activity and identifies subclinical disease. Treg signature Plus mRNA counts by nanoString were normalised, log2-transformed, and split into 50/50 train and test sets before scaling. Utilising our biomarker discovery pipeline, optimum parameters were chosen by elastic net regression on 37 Treg signature Plus genes with leave one out cross validation (LOOCV) for best accuracy in classifying Tregs (sorted CD4+CD25highCD127low) from adult healthy control PB (aHC, signature score assigned towards 0) and active (AJC≥1) JIA PB (signature score assigned towards 1). **A)** Importance of the 23 genes selected in model selection, ranked by frequency of each gene used to differentiate groups in each iteration (importance as %). Genes with 0% importance and not selected for use in final model are not displayed. **B)** Treg signature score of test set, with 0 being most like aHC PB and 1 being most like active JIA PB from model derived through biomarker discovery pipeline. AUC, sensitivity, and specificity shown at cut off 0.5. **C)** Validation of this Treg signature model differentiating paediatric healthy control blood (pHC PB) Tregs from active JIA PB Tregs test sets, with AUC, sensitivity and specificity at cut off 0.5. **D)** Representative heatmap of the mean mRNA counts of 23 genes used in model across aHC, pHC, active JIA and inactive (AJC=0) JIA PB Tregs, clustered by Pearson correlation coefficient with Z-score of expression levels across rows. **E)** Normalised, log2-transformed mRNA counts of CTLA4 and CD96v2 between active and inactive JIA PB Tregs. **F)** Inactive JIA (AJC=0) PB Tregs ran as test dataset on this model, with test aHC and active JIA PB Treg signature scores displayed. clinical Juvenile Arthritis Disease Activity Score (cJADAS) indicated for inactive JIA. Cut-off of 0.5 displayed. **G)** Linear correlation between disease activity (via cJADAS, severe disease >10 removed) and Treg signature score of JIA PB Tregs. 95% confidence bands displayed on linear regression line. **H)** Differentiation between inactive JIA patients with subclinical disease activity (AJC=0, cJADAS≥0.5) and those without (AJC=0, cJADAS<0.5) by Treg signature score with cut-off at 0.5 displayed. For B,C,E,F,H, Mean ±SEM displayed, as well as individual points; cut offs for Sens/Spec calculation displayed by dotted line; Significance determined by Mann-Whitney test (B,C,E,H), and one-way ANOVA with Kruskal-Wallis multiple comparisons post-hoc test (F) with *p<0.05, **p<0.01,****p<0.0001, ns= not significant. PB= peripheral blood; JIA= Juvenile Idiopathic Arthritis; AJC= active joint count; ROC= receiver operating characteristic; AUC= area under curve; Sens= sensitivity; Spec= specificity.

Using this model, with a Treg signature score (0–1) representing the probability of a sample being from an individual with active JIA (AJC≥1, with 1 representing most like active JIA, 0 like aHC), data from inactive (AJC=0) JIA PB Tregs generated scores in a range between aHC PB Tregs and active JIA PB Tregs (Figure 3F). Inactive JIA PB Tregs differentiated significantly from active JIA (mean±SEM inactive 0.4623±0.063 vs active 0.7919±0.043, p=0.0029, Figure 3F), with no overall significant difference from aHC PB Tregs (0.2101±0.027, p=0.2330, Figure 3F). Treg-derived biomarker scores were therefore successful in identifying active disease in the blood of JIA individuals.

To examine the large range of scores from inactive JIA PB Treg samples we explored additional measures of disease activity. cJADAS incorporates AJC with clinician and patient/parent global assessments of disease activity(41). A clinically useful biomarker would need to reflect the level disease activity beyond active joint count, and current biomarkers such as S100 proteins do not correlate with cJADAS (Supplementary Figure S3A). Stratifying Treg signature scores of all JIA PB samples by cJADAS displayed a positive correlation, below a cJADAS of 10 (p<0.0001, r^2^=0.5071, Figure 3G, supplementary Figure S3B), suggesting increased disease activity from low to moderate relates to changes in overall Treg fitness in the blood. Conversely, high disease activity (cJADAS≥10) is likely driven by more than changes in Treg fitness, such as overt inflammation.

Subclinical disease activity, involving symptoms other than inflamed joints such as pain and fatigue, are often noted by patient/parent visual assessment, and difficult to assess objectively in clinic(42). Thus, we further subcategorised the inactive (AJC=0) cohort into remission (cJADAS<0.5) and potential subclinical disease through other recognised symptoms (cJADAS≥0.5), which were significantly different in Treg signature scores (mean±SEM for inactive cJADAS<0.5 0.2848±0.075 vs inactive cJADAS≥0.5 0.6389±0.051, p=0.0111, Figure 3H). Samples without active joint inflammation but with subclinical disease activity were therefore closer to those with active joint flares by Treg signature scores.

With the same cut off of 0.5, defined by aHC vs active JIA PB, the Treg signature scores generated from this blood Treg-derived model objectively distinguished remission from subclinical disease activity without active joint inflammation in JIA (AUC=0.8980) with high sensitivity (0.8571) and specificity (0.8571, Figure 3H).

We previously found that in T1D, a PBMC-derived Treg signature could differentiate between T1D and healthy controls(31, 32). While elastic net regression analysis of 45 genes on unfractionated PBMCs generated a model (supplementary Table S2) to differentiate aHC PBMCs from active JIA PBMCs (AUC=1.000, Supplementary Figure S2A,B), when tested on inactive JIA PBMCs (AJC=0), there was no distinction of inactive from active JIA (AJC≥1), nor subclinical disease activity (AJC=0, cJADAS≥0.5) from remission (Supplementary Figure S2D). This therefore suggests that there is more of an intrinsic change in Treg fitness in active and subclinical disease rather than an overall immunoregulatory imbalance.

We therefore propose a blood Treg-derived model capable of successfully and objectively identifying active and subclinical disease in JIA, recognising Treg fitness as an important factor in disease progression and sustained remission.

### Blood Treg subsets are altered between inactive and active JIA

To investigate whether altered Treg fitness in blood of active JIA could also be assessed by protein expression, we utilised spectral flow cytometry data(19) and unbiased analysis. Many of the genes in the Treg signature could not be assessed for transcribed protein expression by flow cytometry due to inaccessibility or lack of commercial antibody available. However, we have previously developed a 37-parameter spectral panel for comprehensive assessment of cellular composition and phenotype of mononuclear cells, including 14 markers from our Treg signature Plus, which we analysed on PBMCs from individuals with JIA(19). Taking this data for 20 markers associated with Treg function and activation, we performed PhenoGraph cluster analysis(43) on gated live CD3+CD4+Foxp3+ blood Tregs from individuals with clinically active JIA (AJC≥1, n=28) and inactive JIA (AJC=0, n=17 Figure 4A,B).

**Figure 4.**
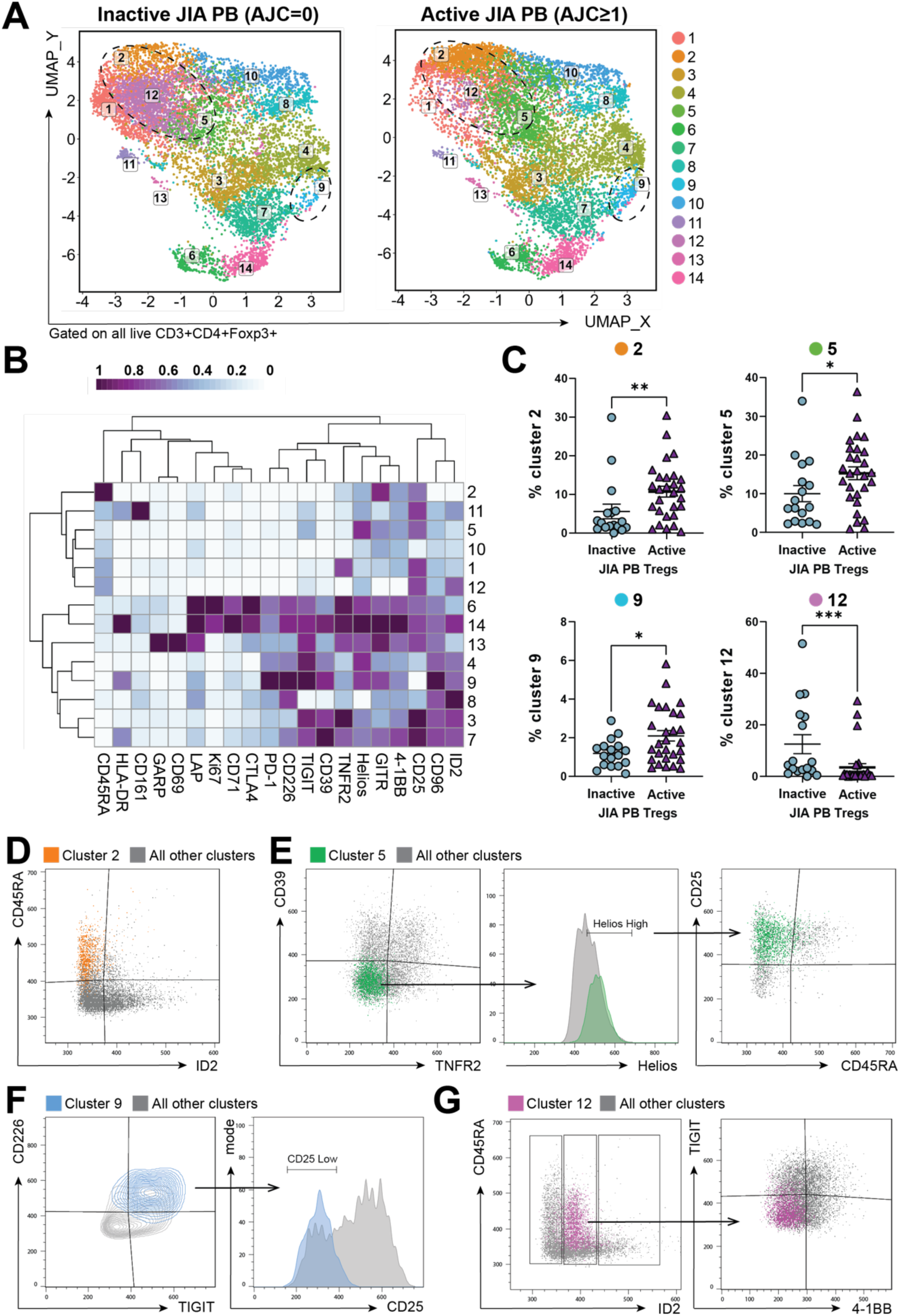
Treg subsets are altered between active and inactive JIA PB. Spectral flow cytometry data analysed from Attrill *et al* dataset(*19*). **A)** UMAP of 14 Treg clusters determined by unbiased PhenoGraph clustering of active (AJC≥1, n=29) and inactive (AJC=0, n=17) PB Tregs, gated on live CD3+CD4+Foxp3+. Clusters which are significantly different in frequency between active and inactive JIA PB Tregs are circled (clusters 2,5,9,12). **B)** Heatmap of 20 markers used for Treg clustering with Z-score for expression levels across cluster rows. **C)** Frequency of clusters 2,5,9, and 12 as a percentage of all Foxp3+ Tregs for individual JIA PB samples with mean±SEM. **D-G)** Representative flow plots displaying phenotypic identities of cluster 2 (D, ID2lowCD45RA+), cluster 5 (E, TNFR2-CD39-HelioshighCD45RAlowCD25high), cluster 9 (F, TIGIT+CD226+CD25low) and cluster 12 (G, ID2intermediate4-1BB-TIGITlow) as coloured overlays to all other JIA PB Treg clusters (grey). Histograms normalised to mode, and flow dot/contour plots display normalised to 100,000 events. Significance determined by Mann-Whitney test (C) where *p<0.05, **p<0.01, ***p<0.001. PB= peripheral blood; JIA= Juvenile Idiopathic Arthritis; AJC= active joint count.

We identified 14 Treg clusters, separated by various combinational protein expression, present in both active and inactive JIA blood, of which four clusters were significantly different between disease activity groups (Figure 4A-C). Clusters 2 (mean±SEM of CD4+Foxp3+ Tregs, inactive 5.59±1.9% vs active 10.71±1.4%, p=0.0051), 5 (inactive 9.99±2.1% vs active 15.24±1.65%, p=0.0352) and 9 (inactive 1.19±0.18% vs active 2.10±0.27%, p=0.0451, Figure 4C) were significantly increased in the blood during an active flare, compared with inactive disease. On the other hand, Treg cluster 12 was predominant in the blood of inactive individuals (12.49±3.7% vs active 3.48±1.4%, p=0.0007, Figure 4C). The identifications of these clusters were validated via manual gating strategies (Figure 4D-G). ‘Active’ Treg clusters were therefore classified as populations of ID2_low_CD45RA+ (cluster 2, Figure 4D), TNFR2-CD39-Helios_high_CD45RA_low_CD25_high_ (cluster 5, Figure 4E), both likely representing resting or possibly latent Treg populations(44, 45), and the effector-like Treg cluster TIGIT+CD226+CD25_low_ (cluster 9, Figure 4F), similar to a Treg subset identified in the inflamed joint(19). The one Treg cluster enriched in patients with clinically inactive JIA was identified to be ID2_intermediate_4-1BB-TIGIT_low_ (cluster 12, Figure 4G), potentially resting Tregs with intermediate levels of ID2, the role of which in Treg maintenance and plasticity is not clear(46, 47).

### Treg cluster ratios correlate with disease activity in JIA and can predict flares

Overall, inactive disease, defined by no active joints, could be most significantly distinguished from active disease by the ratio of identified ‘active’ Treg clusters to that of the Treg cluster associated with inactive disease (frequency of clusters 2+5+9:12, p=0.0001, Figure 5A). Furthermore, this Treg cluster ratio was associated with disease activity (supplementary Figure S3C), capable of significantly differentiating inactive disease (AJC=0, mean±SEM 6.30±2.4) from low disease activity (AJC≥1, 1≤cJADAS<5, 34.9±9.9, p=0.0142) and moderate disease activity (AJC≥1, 5≤cJADAS<10, 66.93±9.7, p<0.0001), as well as low disease activity from moderate (p=0.0164, Figure 5B). High disease activity (AJC≥1, cJADAS≥10, 62.26±25.5), however, did not correlate with Treg cluster ratios, instead appearing to split into two populations (Figure 5B right axis), similar to that seen in Treg gene signature scores (supplementary Figure S3B). This suggests Treg phenotype changes are particularly important in low disease activity range and may be critical for maintaining inactive disease and achieving full remission in JIA.

**Figure 5.**
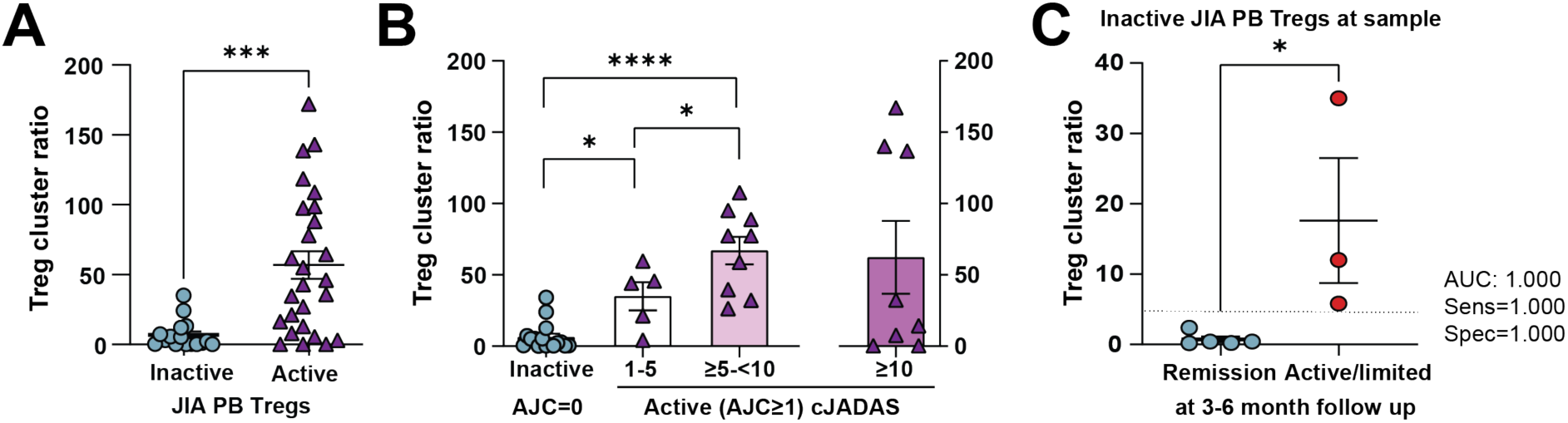
PB Treg cell ratios reflect disease activity and can predict flares in JIA. Ratio of Treg subsets identified by spectral flow cytometry unbiased clustering and classical gating strategies in Fig 4. **A)** Treg cluster ratio of combined Treg clusters [(clusters 2+5+9)/cluster 12] between active (AJC≥1) and inactive JIA (AJC=0) PB Tregs. **B)** Treg cluster ratio across categories of increasing clinical Juvenile Arthritis Disease Activity Score (cJADAS) for active JIA (AJC≥1), with inactive classified as AJC=0. samples from active JIA with ≥10 cJADAS on separate plot to the right. **C)** Treg cluster ratio of inactive samples (AJC=0) at time of sample, classified by disease activity 3-6 months after original sample, where clinical follow-up data was available. Remission without active or limited joints vs presence of active/limited joints recorded at next clinical assessment; ROC curve AUC, Sens and Spec displayed at cut-off of 4.0, shown by dotted line. Significance determined by Mann-Whitney test (A,C), and one-way ANOVA with Krustal-Wallis multiple comparison post-hoc test (B left). *p<0.05, ***p<0.001,****p<0.0001. PB= peripheral blood; JIA= Juvenile Idiopathic Arthritis; AJC= active joint count; ROC= receiver operating characteristic; AUC= area under curve; Sens= sensitivity; Spec= specificity.

Moreover, analysis of measured disease activity at clinical follow up (active/limited joint count 3-6 months after sample was taken) indicates potential predictive power of Treg cluster ratio, with samples with low Treg clusters ratios (<4.0) maintaining remission, distinguished from individuals with higher ratios which went on to flare (p=0.0357, AUC=1.000, Sensitivity= 1.000, Specificity= 1.000 at cut off 4.0, Figure 5C). Therefore, the ratio of Treg cluster frequency could represent a useful biomarker for disease activity to predict flares or indicate full remission, and thus ultimately guide treatment decisions.

### Methotrexate does not affect JIA biomarker potential of Treg fitness measures but alters Treg subsets in blood

We next investigated how medication may affect JIA Treg fitness derived biomarkers using Treg signature scores by mRNA levels and Treg subsets (Treg cluster ratios). Methotrexate (MTX) is widely regarded as a first-choice therapeutic for oligo and RF-polyarticular JIA when non-steroidal anti-inflammatory drugs (NSAIDs) are insufficient(48). To measure the impact MTX has on PB Treg signature scores, and to confirm the difference found between disease activity groups was not due to medication, we divided active and inactive cohorts further into those on MTX at time of sample (MTX) and those not (no MTX, a mix of treatment naïve and those on additional medication). MTX did not significantly alter Treg signature scores within either inactive (AJC=0) or active (AJC≥1) cohorts (Figure 6A), and the distinction between inactive and active patients was still prominent regardless of MTX status (Figure 6A). For Treg subsets defined by protein expression, the Treg cluster ratio (frequency of clusters 2+5+9:12, Figure 4,5) was also still capable of differentiating inactive from active JIA PB on or off MTX (no MTX p=0.0001, MTX p=0.0099, Figure 6B). However, a statistically significant reduction in Treg cluster ratio was found in patients on MTX within both inactive and active groups (inactive no MTX mean±SEM 11.91±4.2 vs inactive MTX 2.01±1.5, p=0.0047; active no MTX 79.03±13.1 vs active MTX 34.84±12.37, p=0.0102, Figure 6C).

**Figure 6.**
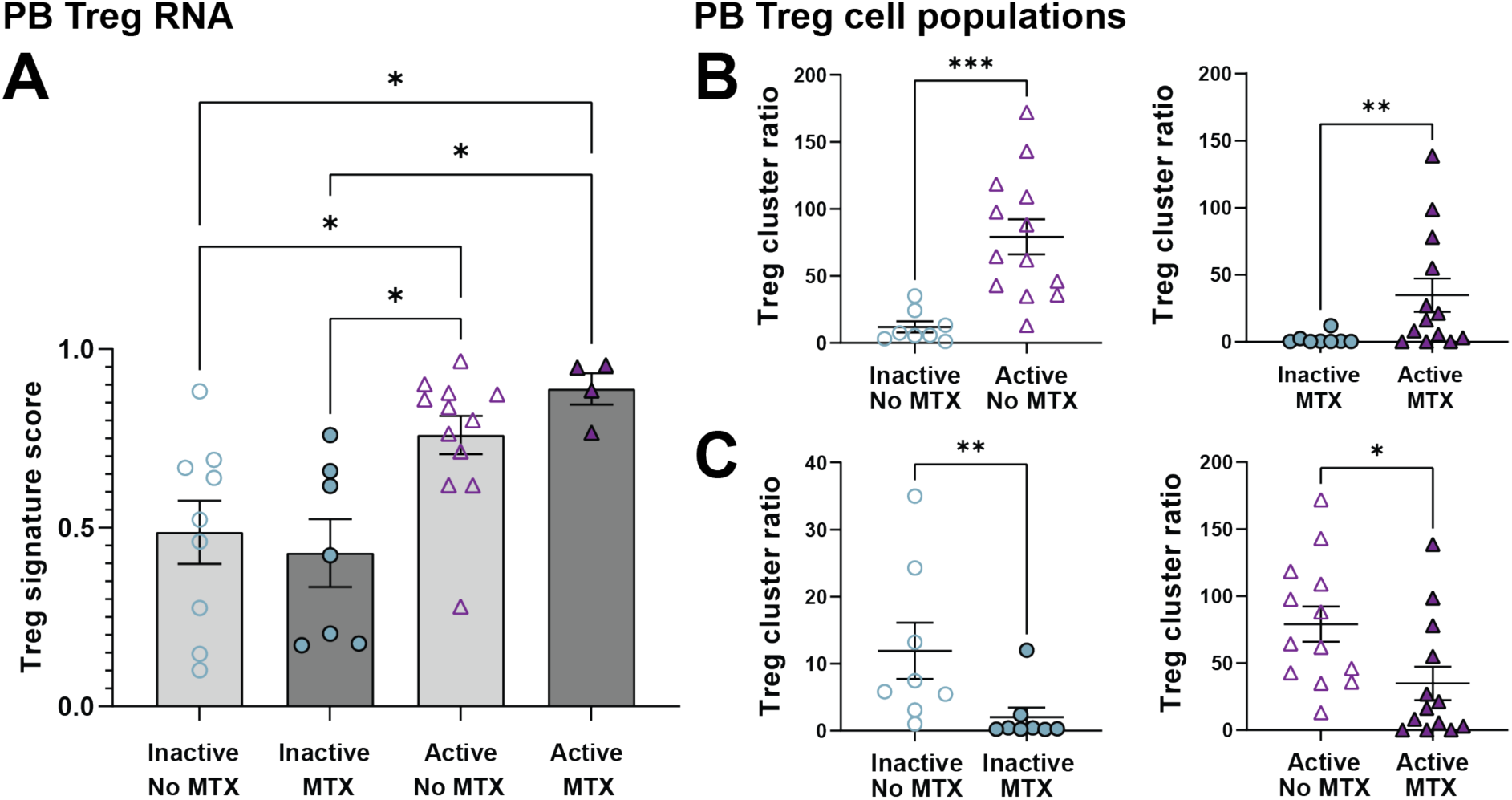
Biomarker potential of Treg fitness measures of disease activity in JIA are unaffected by methotrexate but PB Treg cluster ratio is reduced. Treg fitness measures derived from mRNA nanoString analysis biomarker discovery pipeline (A) or cluster ratios identified via spectral flow cytometry (B-C). **A)** Treg signature scores of JIA PB Treg test sets on (MTX) or off (no MTX) methotrexate at time of sample, determined from 23 gene Treg signature model with 0 being most similar to HC PB Treg and 1 being most similar to active (AJC≥1) JIA PB Treg; Inactive JIA classified as AJC=0. **B)** Treg cluster ratios [(clusters 2+5+9)/cluster 12] for active (AJC≥1) and inactive (AJC=0) JIA PB samples grouped by methotrexate status (off (no MTX, left) and on methotrexate (MTX, right)). **C)** Treg cluster ratios [(clusters 2+5+9)/cluster 12] by methotrexate status grouped by disease activity status (inactive (AJC=0) left and active (AJC≥1) JIA PB right) showing that methotrexate reduces PB Treg cluster ratios within activity groups. Throughout, individual data points with mean±SEM displayed. Significance determined by one-way ANOVA with Krustal-Wallis multiple comparison post-hoc test (A), and Mann-Whitney test (B-C) with *p<0.05, **p<0.01, ***p<0.001, statistical comparisons classed as not significant not displayed. PB= peripheral blood; JIA= Juvenile Idiopathic Arthritis; AJC= active joint count; MTX-= methotrexate

Therefore, unlike previous studies in JIA which suggested MTX does not alter Treg phenotype and function by individual marker expression(49), we propose that MTX may alter specific Treg cluster frequencies towards those reflected in remission, and thus its therapeutic benefit may partially be through changes to Treg fitness. Importantly, the biomarker potential of Treg cluster ratio and Treg signature scores was maintained regardless of MTX status, and thus presents a feasible possibility for clinical translation to guide disease management and treatment decisions.

## Discussion

The importance of Tregs in maintaining tolerance has been well recognised, with various therapeutic approaches now targeting Tregs in an attempt to restore immune homeostasis in autoimmunity(15). In the childhood-onset autoimmune condition JIA, Treg dysfunction has been suggested as a key element of disease pathogenesis, and progression, which is largely unpredictable(1, 16, 17). Here, we investigated the role of Treg fitness in governing disease activity of JIA and its biomarker potential, through the expression of Treg signature genes and proteins that likely correspond to overall functionality.

We implicate Treg fitness to be integral in maintaining JIA remission, utilising mRNA levels of our Treg signature and protein expression with altered Treg subsets. The PB Treg-derived model using our nanoString Treg signature Plus, generated to differentiate healthy controls (HC) from individuals with active JIA, successfully discriminated inactive disease from active, with Treg signature scores of individuals with inactive JIA aligning closer to healthy controls. Further interrogation of the expression of 23 genes governing this model suggests a possible shift towards a more effector-like signature in JIA Tregs, with expression of genes usually upregulated in Tregs compared to effector T cells, such as TRIB1, CSF2RB, ZC2HC1A, HPGD and STAM(31, 50), represented more in a HC Treg signature. The lower expression of these genes in Tregs from across the JIA cohort compared to HC could therefore correspond to diminished regulatory function, with hydroxyprostaglandin dehydrogenase (HPGD) identified as an important tissue-dependent suppressive mechanism of Tregs(51), and CSF2RB (CD131) associated with higher FOXP3 expression(52). However, the access to healthy age-matched controls in this study was limited and additional assessments are required to reveal possible functional outcomes of these Treg signature changes in JIA. Although it was the cumulative differential expression of slight alterations in genes which successfully distinguished active from inactive disease (defined by AJC), CLTA4 and CD96 variant 2 (CD96v2) were statistically different on absolute mRNA count alone. The increased expression of CTLA4, a key immune regulator, on Tregs from individuals with active joints could represent a heightened Treg activation state in the periphery, even when inflammation is localised to the joint(39). Sharing the ligand CD155 with the co-stimulatory CD226 and co-inhibitory TIGIT, there has been conflicting evidence on the role of CD96(53, 54). However, CD96v2 has a higher binding capacity for CD155 than CD96v1(40), and thus might be more potent functionally. The preferential expression of CD96v2 on inactive JIA Tregs, with CD96v1 expression unchanged between inactive and active, could therefore alter functional outcomes of this important co-receptor axis in Tregs.

Interestingly, TIGIT and CD226 protein expression defined an effector-like Treg subset predominant in active JIA PB (cluster 9, TIGIT+CD226+CD25_low_), similar to that seen in the inflamed joint(19). TIGIT is an important co-inhibitory receptor on Tregs, yet the effect of CD226, a co-stimulatory marker, on Treg function has been questioned(55, 56). Additionally, CD25 is crucial for Treg function and lower expression on this cluster may result in reduced proliferation and suppressive capabilities(57). CD25_low_Foxp3+ Tregs have also been proposed as a marker for terminal differentiation into unresponsive Tregs in autoimmunity, failing to regulate overt effector T cell responses(58). Further investigation of this TIGIT+CD226+CD25_low_ cluster seen in active JIA, and the impact of the co-receptor balance in orchestrating Treg fitness, is therefore needed.

From the other three Treg clusters that were identified to be significantly different between inactive and active JIA blood, ID2 was an important distinguishing marker. ID2_low_CD45RA+ was identified as an ‘active’ Treg cluster, whereas the subsets that was most associated with inactive JIA was ID2_intermediate_4-1BB-TIGIT_low_. ID2 is a DNA-binding inhibitor important in T cell maturity and differentiation. In mice, ID2 has been implicated in promoting Treg plasticity into Th17-like effector phenotypes(46), whilst ID2 depletion has also shown inflammatory disease development through lack of Treg maintenance(47). While exact functions of ID2 in Tregs are still unclear, intermediate expression levels of ID2, as found in Treg cluster 12 associated with inactive JIA, could be key in maintaining remission. We therefore propose a change to overall Treg fitness in the blood of active JIA, reflected in our Treg gene signature and altered protein expression. Whether these measurements reflect early Treg response to joint inflammation in the periphery or a driver of active disease remains to be clarified. Further investigation into how the identified genes and proteins influence remission or active disease could present novel therapeutic targets in JIA to sustain remission.

cJADAS is a compound measure of JIA disease activity by including visual global assessment of physicians and patients/parents(41). Both RNA Treg signature scores and Treg cluster ratios (Treg subsets) positively correlated to cJADAS. We therefore offer a quantifiable and objective measure of JIA disease activity which may discriminate low or subclinical levels of disease, for which there is currently no clinically available biomarker. Notably, high disease activity (cJADAS>10) no longer associated with either Treg signature score or Treg cluster ratio, suggesting that severe disease is governed by more than changes in Treg fitness, with overt inflammation likely playing an overriding role at high disease activity. Thus, a possible treatment strategy of first dampening overt inflammation before enhancing Treg fitness could improve outcomes and achieve remission in more children and young people with JIA. One in four patients also receive high patient/parent global visual analogue scale (VAS) scores, despite no active joints and low physician VAS, resulting in higher cJADAS in otherwise clinically inactive individuals(42). Patient/parent VAS is often driven by persistence of morning stiffness, fatigue, and pain, even without clinically apparent inflamed joints(1, 42), likely contributed to by subclinical disease activity which is difficult to measure. Here, we were able to quantify subclinical disease by Treg signature scores and distinguish subclinical disease (AJC=0, cJADAS≥0.5) from full remission (AJC=0, cJADAS<0.5) in otherwise clinically inactive individuals. Furthermore, clinical assessment alone can miss early signs of synovitis in over a third of cases presumed to be inactive(4) and therefore a biological measure to predict flares would aid treatment decisions to sustain remission. Our Treg cluster ratios showed biomarker potential in predicting flares by the next follow up appointment in children/young people that were inactive at time of sample. The predictive capability of Treg signature scores could not be assessed due to limited clinical follow up data for samples analysed by nanoString post-QC. To validate the predictive power of Treg fitness derived measures in JIA, a longitudinal cohort with clinical follow-up data will be needed.

Methotrexate (MTX) is a widely used first-line medication for oligo- and RF-polyarticular JIA(48) and thus, it is important that any biomarker to guide clinical practice can perform regardless of medication status. The biomarker potential in measuring disease activity by Treg signature scores and Treg cluster ratios remained effective independent of MTX status. Moreover, there has been various attempts to measure and predict responders to MTX in JIA(8–11, 59), yet the effect on immune responses is still not fully understood. Despite previous suggestion that MTX does not alter Treg phenotype and function in JIA(49), here, whilst MTX had no effect on Treg signature scores, individuals on MTX at time of sampling showed reduced Treg cluster ratios. This may suggest that MTX, although possibly not altering expression of singular Treg markers, affects Treg subsets in the blood. Similar studies in rheumatoid arthritis have also suggested that MTX can restore suppressive functions of Tregs(60, 61). Therefore, functional investigation of the identified Treg clusters associated with active and inactive disease, and the affect by MTX treatment, may provide insight into MTX mechanism in autoimmunity and Treg cluster ratios could be adapted as a treatment response biomarker to indicate when remission has been achieved.

We therefore establish Treg fitness-based biomarkers in JIA, through Treg signature scores and Treg cluster ratios that can objectively measure, and potentially predict, JIA disease activity. With further prospective and longitudinal validation, Treg fitness-derived biomarkers could become objective clinical tests, showing promise to translate to a valuable tool in clinical practice, requiring only a small blood sample. Monitoring Treg fitness could therefore guide treatment decisions, prevent flares through earlier therapeutic intervention and identify who can safely withdraw from medication without risking an imminent flare. Moreover, Treg-derived biomarkers could also be useful across other autoimmune conditions and thus may have a wider impact on managing autoimmunity and more people achieving long-lasting remission.

## Methods

### Sample collection, processing and demographics

Peripheral blood (PB) was collected via venepuncture from paediatric (<16 years old) healthy individuals (pHC, n=5), adult (≥18 years old) healthy volunteers (aHC, n=26) and a cohort of individuals with diagnosed rheumatoid factor negative (RF-) polyarticular or oligoarticular Juvenile idiopathic arthritis (JIA, n=59). Synovial fluid (SF, n=29) was also collected from unmatched JIA patients via joint aspiration prior to therapeutic intra-articular joint injection. PB and SF was processed as soon as possible for mononuclear cell (MC) isolation via density gradient centrifugation and cryopreservation. Hyaluronidase (1 μL/mL) was additionally added to diluted SF samples, incubated for 30 mins at 37°C, prior to centrifugation.

For JIA patients, available data which was assessed clinically was extracted from study databases or fully anonymised clinical records at time of sample, including disease duration, medication, and clinical Juvenile Arthritis Disease Activity Score (cJADAS), which encompasses active joint count (AJC, /10), physician’s global assessment (/10) and patient/parent global assessment (/10)(41) for PB samples. Inactive JIA was classified as no active joints (AJC=0) at time of sample, with individuals with active JIA classified as having one or more active joints (AJC≥1), via clinical assessment. Table 1 displays sample demographics and clinical characteristics of this cohort.

**Table 1.**
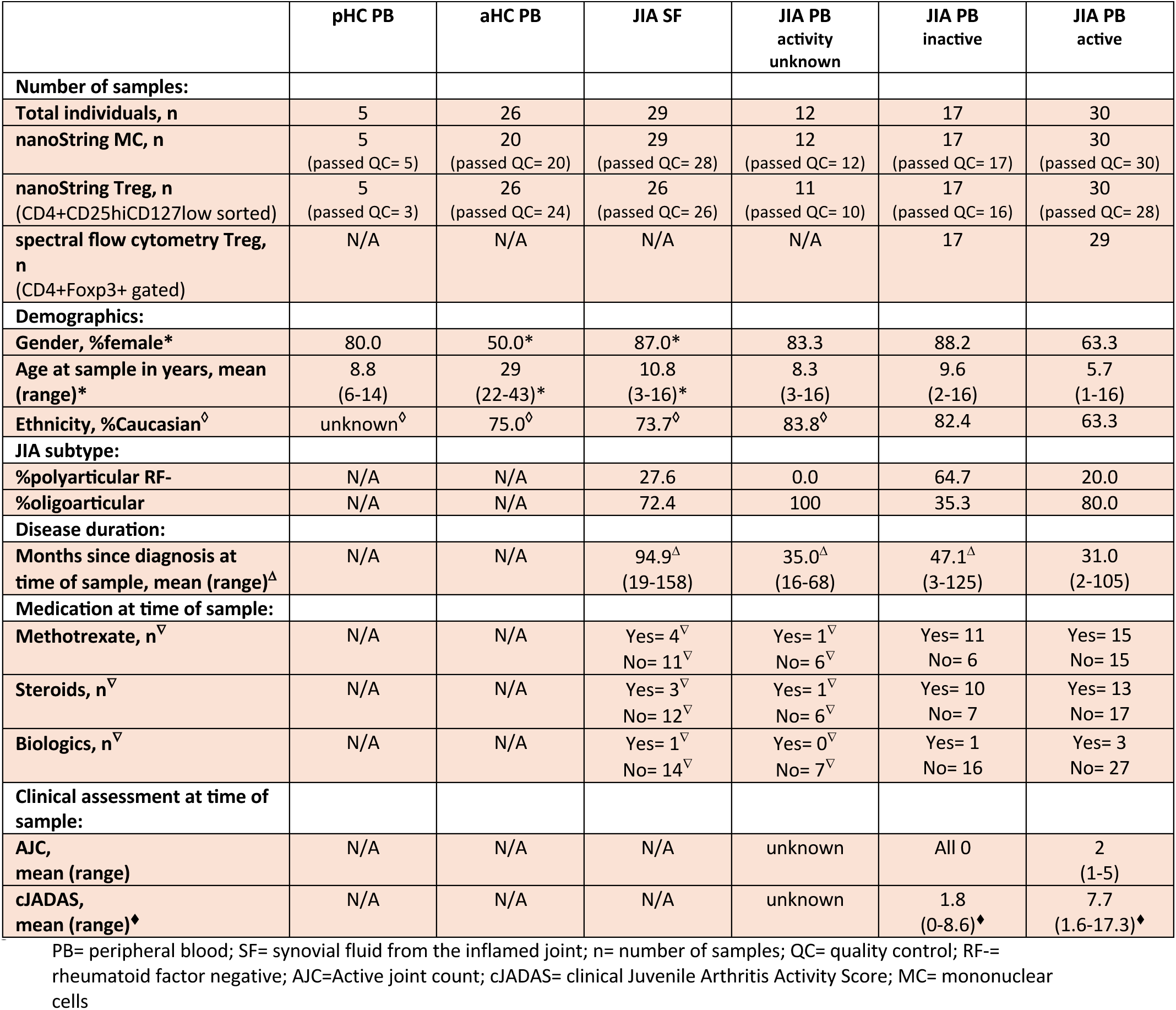
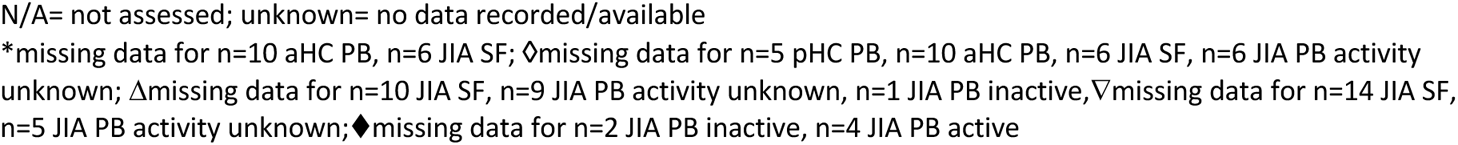
Sample demographics and clinical characteristics of JIA cohort groups.

|  | pHC PB | aHC PB | JIA SF | JIA PB activity unknown | JIA PB inactive | JIA PB active |
| --- | --- | --- | --- | --- | --- | --- |
| **Number of samples:** |  |  |  |  |  |  |
| **Total individuals, n** | 5 | 26 | 29 | 12 | 17 | 30 |
| **nanoString MC, n** | 5<br>(passed QC= 5) | 20<br>(passed QC= 20) | 29<br>(passed QC= 28) | 12<br>(passed QC= 12) | 17<br>(passed QC= 17) | 30<br>(passed QC= 30) |
| **nanoString Treg, n**<br>(CD4+CD25hiCD127low sorted) | 5<br>(passed QC= 3) | 26<br>(passed QC= 24) | 26<br>(passed QC= 26) | 11<br>(passed QC= 10) | 17<br>(passed QC= 16) | 30<br>(passed QC= 28) |
| **spectral flow cytometry Treg, n**<br>(CD4+Foxp3+ gated) | N/A | N/A | N/A | N/A | 17 | 29 |
| **Demographics:** |  |  |  |  |  |  |
| **Gender, %female\*** | 80.0 | 50.0* | 87.0* | 83.3 | 88.2 | 63.3 |
| **Age at sample in years, mean (range)\*** | 8.8<br>(6-14) | 29<br>(22-43)* | 10.8<br>(3-16)* | 8.3<br>(3-16) | 9.6<br>(2-16) | 5.7<br>(1-16) |
| **Ethnicity, %Caucasian<sup>◊</sup>** | unknown<sup>◊</sup> | 75.0<sup>◊</sup> | 73.7<sup>◊</sup> | 83.8<sup>◊</sup> | 82.4 | 63.3 |
| **JIA subtype:** |  |  |  |  |  |  |
| **%polyarticular RF-** | N/A | N/A | 27.6 | 0.0 | 64.7 | 20.0 |
| **%oligoarticular** | N/A | N/A | 72.4 | 100 | 35.3 | 80.0 |
| **Disease duration:** |  |  |  |  |  |  |
| **Months since diagnosis at time of sample, mean (range)<sup>Δ</sup>** | N/A | N/A | 94.9<sup>Δ</sup><br>(19-158) | 35.0<sup>Δ</sup><br>(16-68) | 47.1<sup>Δ</sup><br>(3-125) | 31.0<br>(2-105) |
| **Medication at time of sample:** |  |  |  |  |  |  |
| **Methotrexate, n<sup>▽</sup>** | N/A | N/A | Yes= 4<sup>▽</sup><br>No= 11<sup>▽</sup> | Yes= 1<sup>▽</sup><br>No= 6<sup>▽</sup> | Yes= 11<br>No= 6 | Yes= 15<br>No= 15 |
| **Steroids, n<sup>▽</sup>** | N/A | N/A | Yes= 3<sup>▽</sup><br>No= 12<sup>▽</sup> | Yes= 1<sup>▽</sup><br>No= 6<sup>▽</sup> | Yes= 10<br>No= 7 | Yes= 13<br>No= 17 |
| **Biologics, n<sup>▽</sup>** | N/A | N/A | Yes= 1<sup>▽</sup><br>No= 14<sup>▽</sup> | Yes= 0<sup>▽</sup><br>No= 7<sup>▽</sup> | Yes= 1<br>No= 16 | Yes= 3<br>No= 27 |
| **Clinical assessment at time of sample:** |  |  |  |  |  |  |
| **AJC, mean (range)** | N/A | N/A | N/A | unknown | All 0 | 2<br>(1-5) |
| **cJADAS, mean (range)<sup>♦</sup>** | N/A | N/A | N/A | unknown | 1.8<br>(0-8.6)<sup>♦</sup> | 7.7<br>(1.6-17.3)<sup>♦</sup> |

PB= peripheral blood; SF= synovial fluid from the inflamed joint; n= number of samples; QC= quality control; RF= rheumatoid factor negative; AJC=Active joint count; cJADAS= clinical Juvenile Arthritis Activity Score; MC= mononuclear cells
N/A= not assessed; unknown= no data recorded/available
\*missing data for n=10 aHC PB, n=6 JIA SF; ◊missing data for n=5 pHc PB, n=10 aHC PB, n=6 JIA SF, n=6 JIA PB activity unknown; Δmissing data for n=10 JIA SF, n=9 JIA PB activity unknown, n=1 JIA PB inactive, ∇missing data for n=14 JIA SF, n=5 JIA PB activity unknown; ♦missing data for n=2 JIA PB inactive, n=4 JIA PB active

### Treg isolation and lysate preparation

A minimum of 5×10^4^ cells were kept for mononuclear cell lysis from PBMC and SFMC samples before CD4+ T cells enrichment via magnetic negative selection (EasySep™, StemCell Technologies). Tregs were then sorted from CD4+ T cells on live (FVD-) CD4+CD25highCD127low. Purity checks were performed on a subset of sorted Tregs, staining intracellularly for Foxp3 (Foxp3/transcription factor staining buffer, eBioscience™). Sorted Tregs were lysed, along with mononuclear cell samples, using RNeasy Lysis Buffer (Buffer RLT) with 1% 2-Mercaptoethanol at 1μL/5000 cells. Lysed samples were stored at −80°C until nanoString assessment.

### nanoString

mRNA gene expression was measured for 48 genes from Treg and MC samples using nanoString’s nCounter XT assay, using our custom human Treg signature plus CodeSet (nanoString, CodeSet name Hu_TregsPlus (Pesenacker), supplementary Table S1). Cell lysates were hybridised according to standard protocol, with Reporter CodeSet and Capture ProbeSet, for 18 hours at 65°C before loading onto the Prep station with cartridge and reagent. Loaded cartridges were then transferred to the nCounter Pro digital analyser for mRNA count readouts. A pool of all 48 oligonucleotides at six different concentrations (0 fM-50 fM) was additionally run in triplicate for standard curve and future batch correction, with a 50 fM standard included in each cartridge as a control reference.

### QC and normalisation

Quality control (QC) was performed on nanoString mRNA count data using the R package NanoString quality control dashboard (NACHO)(62), visualising QC parameters (average counts, binding density, median counts) and expression of control genes. Positive control normalisation and total sum normalisation was performed, to a normalisation factor of 5000 counts, with zero counts remaining as zero even after transformation (R scripts will be made available at GitHub Pesenacker lab). Samples which failed QC metrics, positive control normalisation or total sum normalisation, were flagged and removed from further analysis. Data was then log_2_-transformed for input into the biomarker pipeline (R scripts will be made available at GitHub Pesenacker lab).

### Biomarker discovery pipeline

For biomarker discovery, genes with detectable counts (>0) in fewer than 30% of samples across all groups were removed from further analysis to minimise effects driven by outliers (genes removed in each biomarker discovery pipeline shown in supplementary Table S2). In this study, biomarker discovery by elastic net regression was utilised to generate models to differentiate:

1. adult HC PB Tregs vs JIA SF Tregs (SF Treg signature model, input of 42 genes)
2. adult HC PB Tregs vs active (AJC≥1) JIA PB Tregs (Treg signature model, input of 37 genes)
3. adult HC PBMCs vs active (AJC≥1) JIA PBMCs (PBMC Treg signature model, input of 45 genes)

Data from each group were split into equally sized train and test sets. Leave one out cross validation (LOOCV) on the train dataset was used to determine optimal regularisation parameters of the elastic net model (alpha and lambda values, supplementary Table S2) that maximize model accuracy. The importance of different genes was determined in model selection, ranked by frequency of each gene used to differentiate groups in each iteration (importance as %). Gene coefficients not equal to 0 in the final model were determined (supplementary Table S2). Biomarker scores were generated from test datasets using fitted models, and classification performance was assessed via the area under the receiver operating curve (ROC AUC), as well as sensitivity and specificity at a cutoff score value as specified. Separate test sets, including inactive (AJC=0) JIA PB and paediatric HC, were additionally used to further validate the predictive power of the biomarker scores. Details of generated models are provided in supplementary Table S2.

### mRNA expression data visualisation

Principal component analysis (PCA) with normalised, log_2_-transformed data for all 48 genes was conducted via R. Broad Institute public server (Morpheus, https://software.broadinstitute.org/morpheus) was used to generate heat maps of mean mRNA counts of each group of genes in the final model, using Pearson correlation coefficient to cluster closely related gene expression across groups.

### Flow cytometry and clustering analysis

Treg data (live (FVD-) CD3+CD4+Foxp3+) for active (AJC≥1, n=29) and inactive (AJC=0, n=17) JIA PB samples were extracted from the 37-parameter spectral flow cytometry dataset of Attrill *et al*(19), FLOWRepository ID FR-FCM-Z6VC. Unbiased clustering analysis via PhenoGraph was performed on Tregs across 20 markers (CD45RA, HLA-DR, CD161, GARP, CD69, LAP, Ki67, CD71, CTLA4, PD-1, CD226, TIGIT, CD39, TNFR2, Helios, GITR, 4-1BB, CD25, CD96, ID2) using the R package Spectre(43), allowing additional data integration, dimensionality reduction and visualisation. Gating was performed on exported FCS files on FlowJo v10 (BD Biosciences) post-clustering to identify Treg populations and marker expression.

### Statistical analysis

Statistical analysis and data presentation was performed on Graphpad Prism v10.0.3 and R. Mann-Whitney test was performed when comparing two groups, and one-way ANOVA with Kruskal-Wallis multiple comparisons post-hoc test conducted for three or more group comparisons. P values <0.05 were considered significant. Error bars represent standard error of the mean (SEM).

### Study approval

This research was conducted under the informed consent of parents/carers with age-appropriate assent for those under 16 years old, and informed consent for those participants over 16 years according to the Declaration of Helsinki in accordance with the approval of following research ethics committees: NHS London – Bloomsbury/Harrow Research Ethics Committee REC references JIAP-95RU04, CHARMS-05/Q0508/95 studies (Wedderburn) and REC11/LO/0330 (Ciurtin), UCL research Ethics 14017/001 and 14017/002 (Pesenacker).

## Data availability

nanoString data will be made available on Gene Expression Omnibus (https://www.ncbi.nlm.nih.gov/geo/). R scripts for nanoString data normalisation, visualisation and biomarker discovery pipeline will be made available at GitHub Pesenacker lab. Spectral flow cytometry data is available at www.flowrepository.org under FLOWRepository ID FR-FCM-Z6VC.

## Author contributions

MHA contributed to the overall study design, execution of all experiments, data analysis, and writing and editing of the manuscript. DS performed data analysis, figure generation and contributed to writing and editing of the manuscript. TMV, MM helped with data normalisation and biomarker pipeline script development. NMdG and ECR performed a set of Treg sorts; ECR contributed to editing of the manuscript. BJ and MK assisted with sample collection, processing, database management clinical data queries and editing of the manuscript. JIA samples were part of the CHARMS and JIAP studies. LRW oversaw subject recruitment, sample biobanking and patient data collection and contributed to editing of the manuscript. AMP led and oversaw the overall study design, acquisition, analysis and writing and editing of the manuscript, and takes responsibility for the content of the article. All authors read and approved the final version of the manuscript.

## Acknowledgements

This project was funded by NIHR BRC UCLH BRC764/III/AP/101350; and ARUK (now Versus Arthritis) CDF 21738, Versus Arthritis 23159 and 23135, UKRI BBSRC BB/V009524/1 supported salaries of AMP, MHA, DS and contributed to general shared lab consumables. TMV was supported by a Rani Rawji studentship from Shionogi. MM was supported by a Cancer Research UK PhD studentship (A29287). BJ was supported by Fight 4 Sight Versus Arthritis PhD studentship 24VA22. NMdG and ECR were supported by medical research foundation fellowship (MRF-057–0001-RG-ROSS-C0797), with ECR supported by a Kennedy Senior Research fellowship (KENN 21 22 09) and Foreum research career grant (074). The work was also supported by Versus Arthritis 20164, 21593 and 22084, GOSH Children’s Charity (VS0518) and UKRI MRC CLUSTER-JIA award (MR/R013926/1), which supported salaries of MK. The CLUSTER-JIA MRC award provided support for recruitment and sample cohorts; CLUSTER is additionally supported by funding, or funding in kind, from Pfizer, UCB, AbbVie, SOBI and GSK, but those parties played no role in this work. LRW and the Wedderburn lab are supported by the NIHR Biomedical Research Centre (BRC) at Great Ormond Street Hospital.

The authors would like to thank all the patients, parents, clinical staff, and study coordinators at Great Ormond Street Hospital, University College London Hospital and healthy controls for their contribution to this research. We thank Prof Coziana Ciurtin for helpful discussions and for providing access to samples. We acknowledge and thank the contribution of the Centre for Adolescent Rheumatology Versus Arthritis at UCL, ULCH and GOSH, and the CHARMS and JIAP studies at GOSH/UCL GOS ICH and CLUSTER Consortium for providing access to samples and data. We acknowledge and thank Janani Sivakumaran Nguyen at the IIT flow cytometry facility for all her technical help and support. We would like to thank Dr Andreas Mayer and Prof Lucy Walker for helpful discussions and reading of the manuscript. The authors would like to thank the teams at UCL IIT, UCL Rayne Institute and UCL GOS ICH for all the discussion, feedback, and exchange of ideas.

