## supplemental material for "Treg fitness as a biomarker for disease activity in Juvenile Idiopathic Arthritis"

**Supplemental Table S1. Genes in Hu_TregsPlus (Pesenacker) nCounter CodeSet.**
NSID= nanoString internal identifiers**.** Isoforms hit= count of all known transcript variants that probe will hit

| Gene name | Alternative name | Probe NSID | Total Isoforms | Isoforms hit |
|---|---|---|---|---|
| ABCB1 | ABCB1 | NM_000927.3:3910 | 4 | 4 |
| ANK3 | ANK3 | NM_001149.2:1560 | 51 | 51 |
| CD226 | CD226 | NM_001303618.1:844 | 9 | 9 |
| CD96v2 | CD96 | NM_005816.5:543 | 12 | 4 |
| CD96v1 | CD96 | NM_198196.3:544 | 12 | 7 |
| CSF2RB | CSF2RB | NM_000395.2:3300 | 5 | 4 |
| CTLA4 | CTLA4 | NM_005214.3:405 | 2 | 2 |
| DACT1 | DACT1 | NM_001079520.1:3350 | 7 | 7 |
| PMS2L11 | DTX2P1-UPK3BP1-PMS2P11 | NR_023383.1:813 | 1 | 1 |
| EIF3S6 | EIF3E | NM_001568.2:370 | 2 | 2 |
| FOXP3 | FOXP3 | NM_014009.3:1230 | 4 | 4 |
| HDGFRP3 | HDGFL3 | NM_016073.2:639 | 3 | 0 |
| HNRPA1 | HNRNPA1 | NM_002136.2:20 | 3 | 3 |
| HPGD | HPGD | NM_001145816.1:570 | 8 | 4 |
| ICA1 | ICA1 | NM_001136020.1:145 | 42 | 22 |
| ID2 | ID2 | NM_002166.4:505 | 1 | 1 |
| IKZF2 | IKZF2 | NM_016260.2:870 | 27 | 23 |
| IL1R1 | IL1R1 | NM_000877.2:4295 | 17 | 16 |
| IL1R2 | IL1R2 | NM_004633.3:1305 | 14 | 11 |
| IL1RN | IL1RN | NM_173842.1:110 | 7 | 1 |
| IL7R | IL7R | NM_002185.2:1610 | 3 | 3 |
| KLRB1 | KLRB1 | NM_002258.2:85 | 1 | 1 |
| LPIN2 | LPIN2 | NM_014646.2:2170 | 7 | 6 |
| LRRC32 | LRRC32 | NM_005512.2:3470 | 9 | 5 |
| METTL7A | METTL7A | NM_014033.3:280 | 1 | 1 |
| nectin2 | NECTIN2 | NM_001042724.1:1120 | 2 | 2 |
| NELL2 | NELL2 | NM_006159.1:180 | 11 | 11 |
| NSUN5B | NSUN5P1 | NR_033322.2:559 | 2 | 2 |
| PTPRK | PTPRK | NM_001135648.1:4315 | 20 | 16 |
| PVR | PVR | NM_006505.3:604 | 4 | 4 |
| RBMS3 | RBMS3 | NM_001003792.2:210 | 16 | 14 |
| RPL23A | RPL23A | NM_000984.5:611 | 1 | 1 |
| TMEM23 | SGMS1 | NM_147156.3:280 | 7 | 1 |
| STAM | STAM | NM_003473.3:1805 | 11 | 10 |
| TGFB1 | TGFB1 | NM_000660.3:1260 | 2 | 2 |
| TIGIT | TIGIT | NM_173799.2:1968 | 3 | 3 |
| TNF | TNF | NM_000594.2:1010 | 1 | 1 |
| TNFAIP3 | TNFAIP3 | NM_006290.2:260 | 9 | 9 |
| TNFRSF1A | TNFRSF1A | NM_001065.2:515 | 4 | 4 |
| TNFRSF1B | TNFRSF1B | NM_001066.2:835 | 6 | 6 |
| TNFRSF9 | TNFRSF9 | NM_001561.4:255 | 2 | 2 |
| TRIB1 | TRIB1 | NM_025195.2:3460 | 3 | 3 |
| UFC1 | UFC1 | NM_016406.3:380 | 4 | 4 |
| VAV3 | VAV3 | NM_001079874.1:352 | 10 | 9 |
| ECOP | VOPP1 | NM_030796.3:2090 | 17 | 13 |
| ZBTB38 | ZBTB38 | NM_001080412.2:2310 | 86 | 86 |
| C8ORF70 | ZC2HC1A | NM_016010.2:665 | 4 | 4 |
| ZNF532 | ZNF532 | NM_018181.4:875 | 42 | 42 |

Isoforms Not Hit By Probe:
CD96v2 (XR_241462.1;XR_924090.1;XM_017005522.1;XR_001739977.1;XM_017005521.1;XM_005247063.3;XM_006713469.3;NM_198196.3); CD96v1 (XM_006713470.3;XM_017005522.1;NM_005816.5;NM_001318889.2;NR_134917.2); CSF2RB (XM_011529905.2); HDGFRP3 (XR_001751298.2;XM_006720554.4); HPGD (XR_938728.2;NM_001363574.2;NM_001256306.2;NM_001256305.2); ICA1 (XM_011515354.1;XM_011515351.1;XM_011515353.2;XM_017012116.1;XM_011515357.2;XM_024446741.1;XM_011515356.3;XM_011515355.3;NM_001350829.2;NM_001276478.2;NM_001350828.2;NM_001350821.2;NR_146928.2;NM_001350830.2;NM_001350824.2;NM_001350834.2;NR_146926.2;NM_001350823.2;NM_001350836.2;NM_001350825.2); IKZF2 (XM_017003592.2;XM_011510817.3;XM_011510819.3;NM_001371277.1); IL1R1 (NM_001320986.2); IL1R2 (XM_011511807.1;XR_923024.2;NM_001261419.2); IL1RN (XM_011511121.1;NM_173843.3;NM_000577.5;NM_001318914.2;NM_001379360.1;NM_173841.3); LPIN2 (XR_935074.2); LRRC32 (NM_001370191.1;NM_001370187.1;NM_001370188.1;NR_163259.1); PTPRK (XM_011536021.3;XM_011536020.3;NM_001291982.2;NM_001291983.2); RBMS3 (XM_005265065.5;XM_005265063.2); TMEM23 (XM_005269675.1;XM_011539583.2;XR_945651.3;XR_945650.3;XM_011539584.3;XR_001747078.2); STAM (XM_011519695.3); VAV3 (XM_005270361.1); ECOP (XM_011515539.1;XM_011515546.2;XM_011515540.2;XM_011515541.2).

**Supplemental Table S2. Models generated via the nanoString biomarker discovery pipeline.**
Model optimum parameters (opT) chosen from elastic net regression with leave one out cross validation (LOOCV) for best accuracy (RMSE) in classifying control (biomarker score 0) and Juvenile Idiopathic Arthritis (JIA) measure (biomarker score 1) from 48 gene Treg signature Plus nanoString codeSet. The intercept is represented at the start of each model, with coefficients for each corresponding mRNA count. p represents the biomarker score, the probability of a sample being most similar to the JIA measure. aHC= adult healthy control; PB= peripheral blood; SF= synovial fluid; AJC= active joint count.

| | **Control (0)** | **Measure (1)** | **Model opT** | **Genes inputted** | **Genes selected** | **Biomarker score model ln(p/(1-p))=** |
|---|---|---|---|---|---|---|
| 1 | aHC PB Treg | JIA SF Treg | α = 0<br>λ= 1 | n=42<br>(Genes removed for lack of counts across groups: ABCB1, PTPRK, PVR, nectin2, HDGFRP3, NSUN5B) | n=42 | **-0.1763041** +0.064000872(**IL1R1**) -0.061154754(**EIF3S6**) +0.060280298(**CTLA4**) -0.05578962(**HNRPA1**) -0.054019342(**RPL23A**) +0.053685732(**FOXP3**) +0.053680697(**TIGIT**) +0.053352253(**TNFRSF1B**) +0.050697622(**HPGD**) -0.045305873(**ZNF532**) +0.04370134(**IL1RN**) -0.041425091(**RBMS3**) -0.04003701(**KLRB1**) +0.038695421(**TNFRSF9**) -0.038240175(**STAM**) +0.037008964(**ZC2HC1A**) -0.034392396(**TNFAIP3**) +0.033761464(**IL1R2**) -0.033688476(**TMEM23**) +0.031994581(**CD226**) +0.030579686(**UFC1**) +0.029999166(**METTL7A**) +0.023638991(**LPIN2**) -0.023105339(**IL7R**) +0.022636694(**CSF2RB**) +0.018290016(**ZBTB38**) +0.017926852(**ECOP**) -0.017704245(**VAV3**) +0.017467846(**CD96v1**) +0.016955562(**IKZF2**) -0.016119258(**NELL2**) +0.015771323(**TRIB1**) +0.015703359(**TNFRSF1A**) -0.015252279(**ANK3**) -0.011676661(**DACT1**) +0.007778844(**PMS2L11**) +0.007709801(**ICA1**) +0.007628293(**TGFB1**) +0.005649417(**TNF**) -0.00546917(**ID2**) -0.003547315(**LRRC32**) -0.002807969(**CD96v2**) |
| 2 | aHC PB Treg | Active (AJC≥1) JIA PB Treg | α = 0.1<br>λ= 0.28 | n=37<br>(Genes removed for lack of counts across groups: ABCB1, PTPRK, PVR, nectin2, HDGFRP3, NSUN5B, IL1R1, IL1RN, IL1R2, ANK3, LRRC32) | n=23 | **-0.212523393** -0.40164806(**ZNF532**) -0.245803195(**TNFRSF9**) + 0.237943766(**IL7R**) -0.214083179(**TNF**) -0.203055138(**ZC2HC1A**) + 0.198345626(**RBMS3**) + 0.18951266(**HNRPA1**) -0.176118497(**STAM**) -0.172881361(**CSF2RB**) -0.171806539(**TRIB1**) -0.153817412(**NELL2**) + 0.131290093(**LPIN2**) -0.114750547(**UFC1**) -0.101507564(**CD96v2**) -0.06832873(**PMS2L11**) + 0.06627762(**TNFRSF1A**) -0.055405803(**CTLA4**) + 0.050597526(**EIF3S6**) -0.046719775(**HPGD**) + 0.039129694(**TNFRSF1B**) -0.033808473(**TNFAIP3**) -0.024183641(**TGFB1**) + 0.012989037(**IKZF2**) |
| 3 | aHC PBMC | Active (AJC≥1) JIA PBMC | α =0.4<br>λ= 0.37 | n= 45<br>(Genes removed for lack of counts across groups: IL1R1, IL1R2, LRRC32) | n=6 | **0.00277288** -0.3989582(**TNF**) +0.23199649(**HNRPA1**) +0.16918882(**DACT1**) -0.1307635(**IL1RN**) -0.1106903(**TNFRSF9**) +0.03880683(**RBMS3**) |

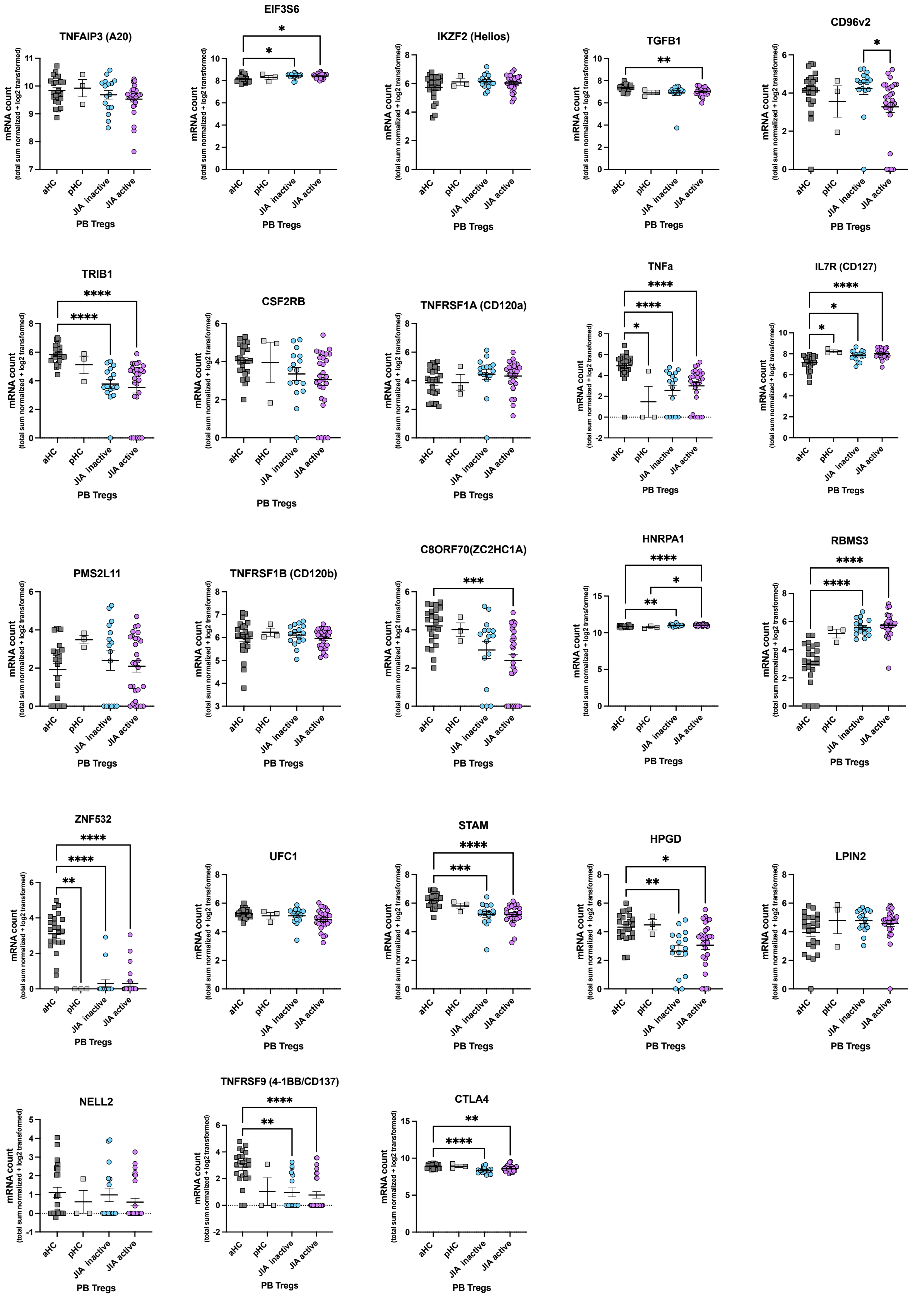


**Supplementary Figure S1. mRNA counts of 23 genes in JIA Treg signature score model.**
mRNA count determined by nanoString Treg signature Plus custom CodeSet on Treg cell lysates, isolated by cell sorting on CD4+CD25highCD127low. Normalised, $log_2$-transformed counts displayed. Significance determined by one-way ANOVA with Kruskal-Wallis multiple comparisons post hoc test. *p<0.05, **p<0.01, ***p<0.001,****p<0.0001. non-significant p values not displayed. PB= peripheral blood; aHC= adult healthy control; pHC= paediatric healthy control; JIA inactive classified as active joint count (AJC)=0, JIA active classified as AJC≥1.

### Model 3: derived from aHC PBMC vs Active JIA PBMC

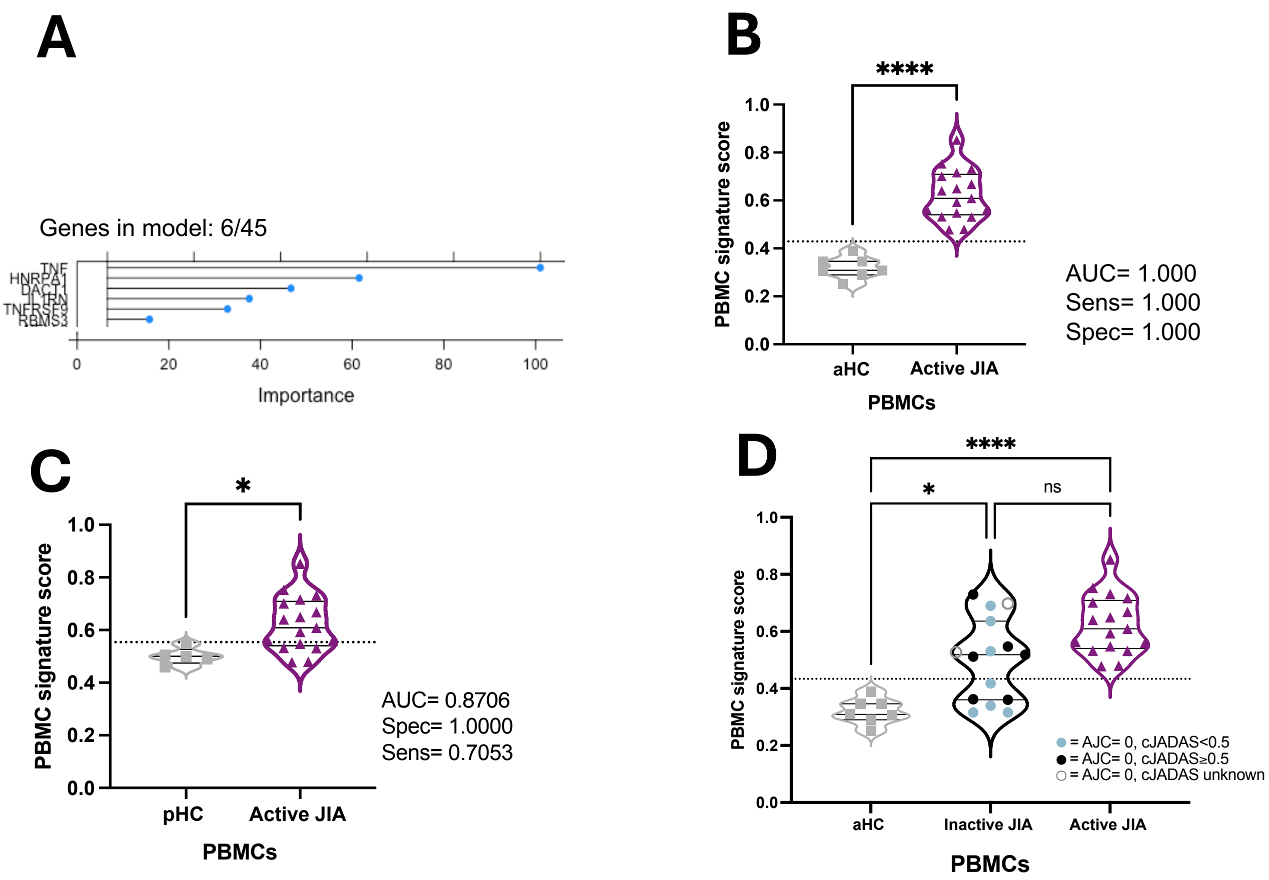


**Supplemental Figure S2. PBMC-derived Treg signature scores in JIA.** Optimum parameters were chosen by elastic net regression on 45 genes with leave one out cross validation (LOOCV) to classify PBMCs from adult healthy control (aHC, PBMC signature score towards 0) and active (AJC≥1) JIA (PBMC signature score towards 1). **(A)** 6 genes selected in the final model are displayed with importance level during LOOCV. **(B)** Datasets were split into unsupervised 50/50 train and test sets, with PBMC signature scores of test sets displayed. AUC, sensitivity and specificity of ROC curve displayed, cut off determined as 0.4335. **C)** paediatric HC (pHC) PBMC test set signature scores in comparison to active JIA PBMC. AUC, sensitivity and specificity of ROC curve displayed, cut off determined as 0.5539. Mann-Whitney test performed (B-C), *p<0.05, ****=p<0.0001. **(D)** Inactive JIA (AJC=0) ran as additional test dataset. ANOVA with Kruskal-Wallis post hoc test performed, *=p<0.05, ****p<0.0001, ns= not significant. PBMC= peripheral blood monocular cell; JIA= Juvenile Idiopathic Arthritis; cJADAS= clinical Juvenile Arthritis Disease Activity Score, comprising of active joint count (AJC, /10), physician's global assessment (/10) and patient/parent global assessment (/10). Sens= Sensitivity. Spec= Specificity. AJC= Active joint count.

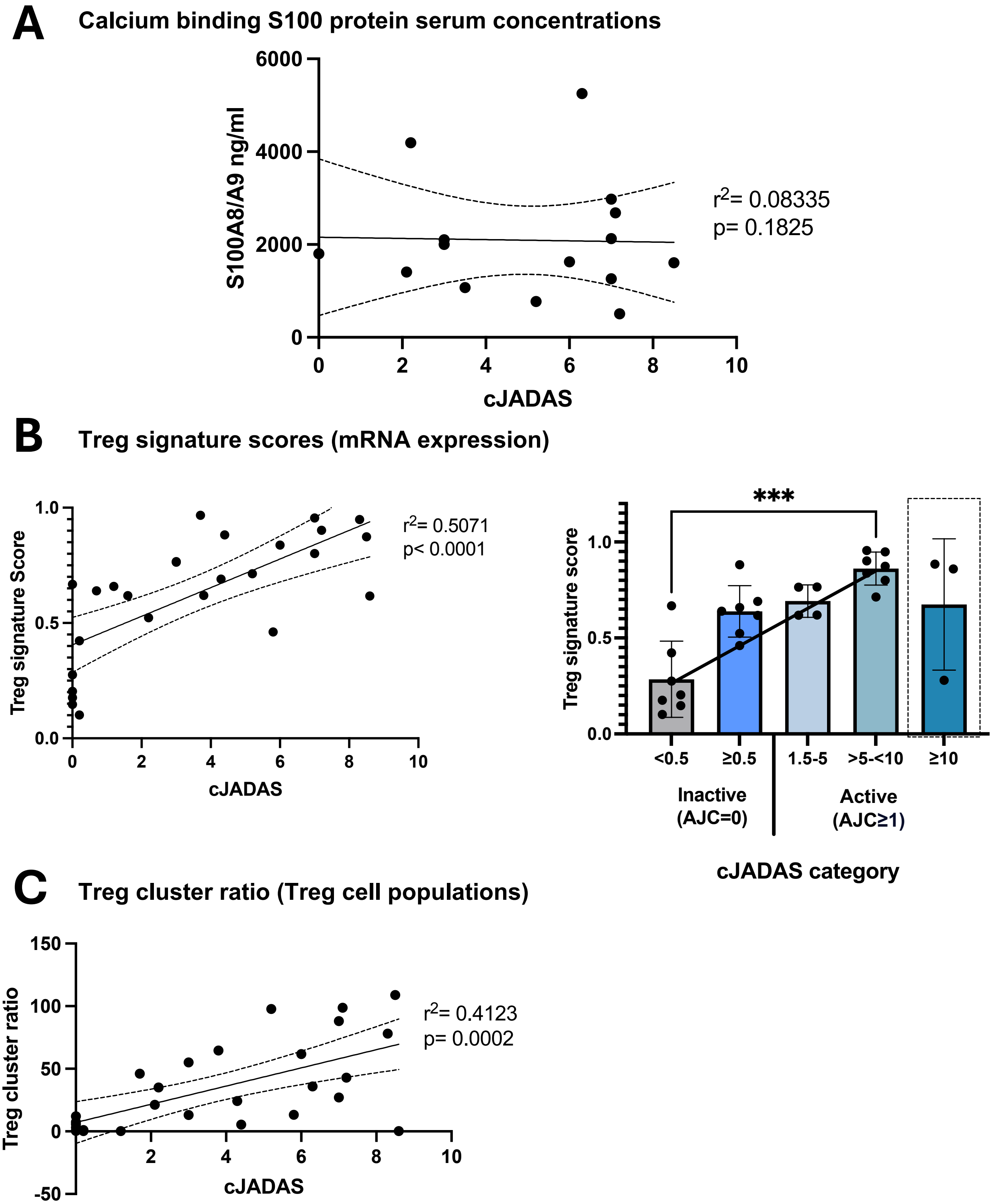


**Supplementary Figure S3. PB Treg fitness-derived measures offer biomarker potential to measure disease activity in JIA that is not currently reflected in clinically available serum markers.** Paired (where available) PB samples of **A)** S100A8/9 serum protein blood concentration, **B)** PB Treg signature scores and **C)** PB Treg cluster ratios against cJADAS. cJADAS>10 not displayed on regression plots. **B)** right, cJADAS by subcategory, ***p<0.001 by one-way ANOVA with Kruskal-Wallis multiple post hoc test. Treg signature score (mRNA) measured from a model of 23 genes to differentiate healthy control from active (AJC≥1) JIA blood Tregs. Treg cluster ratio (subsets via protein expression) consists of the frequency of three Treg clusters associated with active disease/ one Treg subpopulation associated with inactive disease. cJADAS= clinical Juvenile Arthritis Disease Activity Score, comprising of active joint count (AJC, /10), physician's global assessment (/10) and patient/parent global assessment (/10). PB= peripheral blood.
